# Cryo-EM structure of an amyloid fibril from systemic amyloidosis

**DOI:** 10.1101/357129

**Authors:** Falk Liberta, Sarah Loerch, Matthies Rennegarbe, Angelika Schierhorn, Per Westermark, Gunilla T. Westermark, Nikolaus Grigorieff, Marcus Fändrich, Matthias Schmidt

## Abstract

Systemic AA amyloidosis is a worldwide occurring disease of humans and animals that arises from the misfolding of serum amyloid A protein. To provide insights into the molecular basis of this disease we used electron cryo-microscopy and determined the structure of an *ex vivo* amyloid fibril purified from AA amyloidotic mice at 3.0 Å resolution. The fibril consists of C-terminally truncated serum amyloid A protein arranged into a compactly folded all-β conformation. The structure identifies the protein N-terminus as central for the assembly of this fibril and provides a mechanism for its prion-like replication. Our data further explain how amino acid substitutions within the tightly packed fibril core can lead to amyloid resistance *in vivo*.

## Main Text

Systemic AA amyloidosis is a major form of systemic amyloidosis that arises from the formation of amyloid fibrils from serum amyloid A (SAA) protein (*1–3*). The massive deposition of these fibrils in organs like spleen, liver and kidneys physically distorts and impairs the affected tissues (*2, 4*). In addition, there can be toxic effects of oligomeric fibrillation intermediates, similar to other amyloid diseases (*5*). Current treatments, which aim to reduce the level of SAA in the blood, are unable to control the disease in all cases, but therapies that directly target SAA misfolding are not available (*3*). Systemic AA amyloidosis has been a key model for understanding pathological protein misfolding, and its existence in animals has given rise to uniquely relevant mouse and other animal models (*6*). Murine AA amyloidosis reproduces crucial features of the disease process in humans (*2, 7, 8*) and the fibril precursor proteins (mSAA1.1 in mouse and hSAA1.1 and hSAA1.3 in humans) are largely conserved (*9*).

The disease show prion-like characteristics in mice and other animals (*8, 10*). Injection of AA amyloid fibrils or oligomers into mice with high serum mSAA1.1 levels leads to a very fast formation of amyloid deposits in the recipients (*8, 11*). Transmission of the disease to mice is possible via oral uptake and with amyloid fibrils purified from different species, including humans (*6*). However, there is no evidence that human AA amyloidosis is transmissible or contagious. While AA amyloid fibrils are crucial for the prion-like features (*8*) and disease pathology (*2*), their detailed structure remains unknown. In this study, we have used electron cryo-microscopy (cryo-EM) to determine the structure of an AA amyloid fibril from diseased murine spleen.

The protocol to extract the fibrils from amyloidotic tissue avoids harsh physical or chemical conditions and maintains the intact linear architecture of the amyloid fibrils (*12*). Based on transmission electron microscopy (TEM) more than 90 % of the extracted fibrils belong to the same morphology that is defined by a width of ˜12 nm and a cross-over distance of ˜75 nm (fig. S1, A and B). The extracted AA amyloid fibrils contrast morphologically with ‘amyloid-like fibrils’ prepared from full-length mSAA1.1 or the fragment mSAA1.1(1-76) *in vitro* (fig. S1C). AA amyloid fibrils are also more proteinase K resistant than their amyloid-like counterparts and produce a different proteolytic fragmentation pattern (fig. S1, D-F).

Denaturing gel electrophoresis and mass spectrometry (MS) show that the extracted fibrils are free from major protein contaminants (fig. S2, A and B). Consistent with previous observations (*8*), they contain C-terminally truncated mSAA1.1 protein, mainly the fragments mSAA1.1(1-76), mSAA1.1(1-82) and mSAA1.1(1-83) (table S1). Proteinase K treatment removes the two longer fragments, while fibril morphology (*13*), fragment mSAA1.1(1-76) (fig. S2B) and prion-like characteristics are retained. Histological analysis of murine spleen sections with Congo red green birefringence, the gold standard of amyloid detection in medical diagnosis (*1*), shows large amyloid deposits (quantified as amyloid grade +3) in mice that were injected with AA amyloid fibrils, irrespective of proteinase K treatment (fig. S2C). Control animals that received non-fibrillar recombinant SAA1.1(1-76) or a 1:1 mixture of SAA1.1(1-76) and SAA1.1(1-82) (fig. S2C) did not develop amyloid deposits (amyloid grade 0). We conclude that the prion-like activity is associated with a protease resistant conformation adopted by residues 1-76.

Using cryo-EM, we determined the three-dimensional (3D) structure of the major amyloid fibril morphology (no proteinase K treatment) at a resolution of 3.0 Å (Fig. 1; fig. S3A and table S2). Our 3D map resolves most side-chains and allowed us to unambiguously trace and assign residues 1-69 of mSAA1.1 (Fig. 1D and table S3). The density corresponding to residues 70-82 is diffuse (fig. S3B), likely due to structural disorder. Two-dimensional (2D) projections of the fitted model correlate well with the 2D class averages (fig. S3C). The fibril is polar and exhibits pseudo-2^1^ symmetry, resembling previously reported cross-β fibrils (*14–16*). It consists of two identical protofilaments (PFs) that are arranged in parallel. Consistent with the 2D class averages (fig. S3C), they are offset from one another by approximately half a cross-β repeat. The two-PF double helix is left-handed, as confirmed by platinum side shadowing and TEM (fig. S4A).

**Fig. 1.**
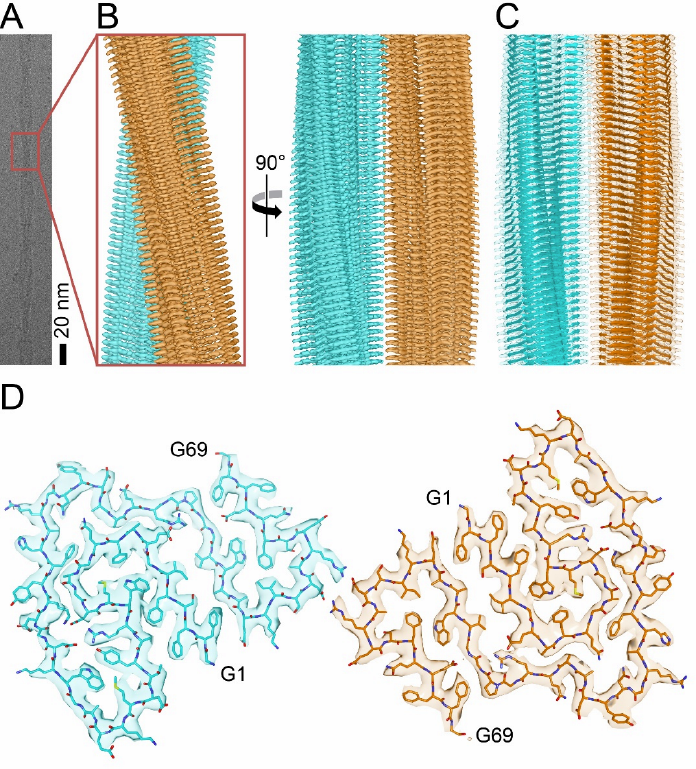
Cryo-EM reconstruction of a murine AA amyloid fibril. (A) Cryo-EM image. (B) Side views of the reconstruction. The two PFs are colored cyan and orange. (C) Model of residues 1-69, superimposed with the density. (D) Cross-sectional view of one molecular layer. The density is superimposed with a molecular model.

Each PF represents a stack of fibril proteins and encompasses nine parallel cross-β sheets (β1-β9) (Fig. 2A), consistent with the general cross-β architecture of amyloid fibrils (*17, 18*). The fibril conformation differs substantially from all known native conformations of SAA family members that uniformly belong to the all-α class of protein folds (Fig. 2B). The conformational differences between the fibril protein and its native precursor resemble the differences between cellular and pathogenic prion protein (*19*) and imply the unfolding of native structure as a prerequisite of fibril formation *in vivo*. The PFs and the polypeptide chains appear slightly tilted relative to the main fibril axis (Fig. 1B) and β-sheets exhibit a left-hand twist when viewed along the fibril axis (fig. S4B). The orientation of the fibril twist corresponds to the β-sheet twist in globular proteins (fig. S4C). The compact fold of the fibril protein and the absence of large cavities (Fig. 1D) are in agreement with the observed proteolytic resistance (fig. S2B). The fibril protein conformation superficially resembles the Greek key topology, but similar to α-synuclein-derived amyloid-like fibrils (*16, 20*), there are no intramolecular backbone hydrogen bonds between the β-strands of the same polypeptide chain as required for a Greek key. Instead, the intramolecular strand-strand interactions are formed by the amino acid side-chains, and we hereafter refer to this motif as an ‘amyloid key‘.

**Fig. 2.**
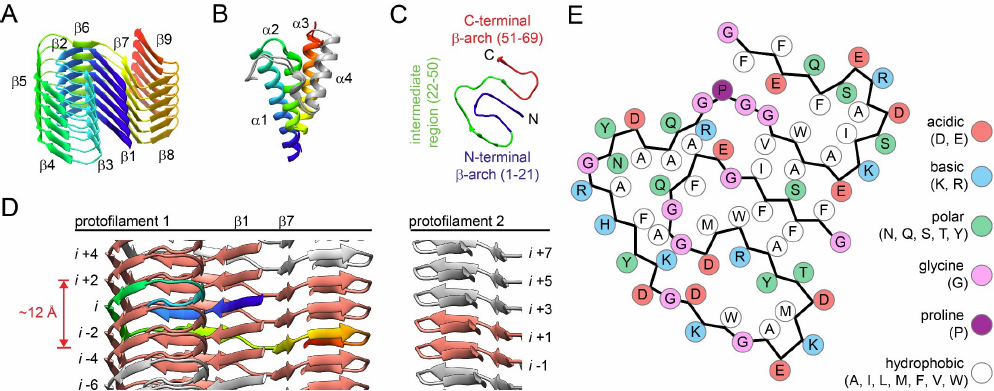
The conformation of the fibril protein. (A) Six molecular layers of a PF. (B) Superposition of the native structure of mSAA3 (PDB 4Q5G) and hSAA1.1 (PDB 4IP8), demonstrating their very close similarity. Residues 1-70 are rainbow-colored from N (blue) to C (red). Grey: C-terminal residues not resolved or missing in the fibril structure. (C) Cross-sectional view of one fibril protein. (D) Fibril side view. Molecule *i* (rainbow-colored) interacts with six other molecules (salmon-colored). The 12 Å chain rise is measured between the carbonyl carbons of Val51 and Lys24. (E) Schematic view of the fibril protein illustrating the complementary packing.

The central structural element of the amyloid key is an N-terminal β-arch (residues 1-21), the most hydrophobic and amyloidogenic segment of the protein (*21–23*). This β-arch is surrounded by the intermediate region (residues 22-50) and the C-terminal β-arch (residues 51-69). N- and C-terminal β-arch are connected by an intermolecular zipper formed by strand β7 of molecule *i* and strands β1 of molecules *i-*2 and *i–*4 (Fig. 2, C and D). Each fibril protein spans ˜12 Å along the fibril axis and interacts with six other protein molecules, four within the same PF and two of the opposite PF (Fig. 2D). This non-planarity originates from the tilt of the PF with respect to the fibril axis (Fig. 1B) and a GPGG motif (residues 47-50) that induces a ˜5.5 Å height change of the polypeptide chain relative to the fibril axis.

The fibril surface is predominantly hydrophilic, whereas residues Phe3, Phe5, Phe10, Met16, Trp17 and Ala19 form a tightly packed hydrophobic core within the N-terminal β-arch (Fig. 2E). The more C-terminal segments of the protein pack onto the outer face of the N-terminal β-arch, mediated by a complex pattern of ionic, polar and hydrophobic interactions (fig. S5). Some of these interactions occur across different molecular layers of the fibril, interdigitating the PF structure. Salt bridges structure the interface between the two PFs which is remarkably small and formed by only two residues (Asp59 and Arg61) from the C-terminal β-arch (Fig. 3). These residues make bidentate, reciprocal contacts with the respective residues in chains *i*+1 and *i*-1 in the opposing PF, which is consistent with the observed pseudo-21 symmetry.

**Fig. 3.**
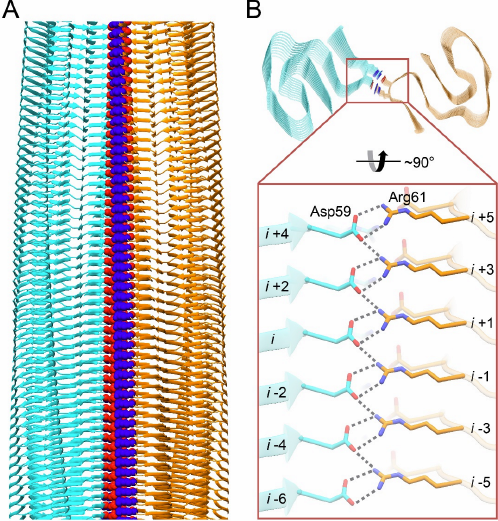
PF interface. (A) Fibril side view. Residues Asp59 (red) and Arg61 (blue) are shown as spacefilling models. (B) Top view and side view close-up of the side view showing the ladder of ionic residues which make bidentate interactions with the other PF.

Our structure leads to a simple mechanism for the protein-like replication of the fibril structure that depends on the β1/β7 zipper region and the exposure of unpaired half zippers at either end of the fibril (fig. S6). Pairing of the half zipper with a complementary strand from an incoming mSAA1.1 molecule helps to orient the newly added chain at the fibril tip and nucleates the formation of a new half zipper in the added molecule such that the fibril structure can be further replicated. A related mechanism has recently been proposed for amyloid-like fibrils formed from Aβ(1-42) peptide (*15*).

The buried position and conformation of residues 1-21 explains the previously described role of the protein N-terminus for fibril formation (*22*) and for the resistance of *Mus musculus czech* and CE/J mice to development of systemic AA amyloidosis. This resistance originates from the expression of mSAA1.5 and mSAA2.2 proteins in these animals (*24, 25*). Both proteins differ from mSAA1.1 by a synonymous Ile6Val substitution, which is consistent with our amyloid fibril morphology and a Gly7His substitution (Fig. 4A) that is structurally disruptive and places a bulky, charged residue into the tightly packed, hydrophobic core of the N-terminal β-arch (Fig. 4B). Importantly, mSAA2.2 is able to readily form amyloid-like fibrils *in vitro* (*23*), demonstrating that the amyloid resistance of CE/J mice does not arise from an intrinsic inability to form cross-β structures but rather from its specific incompatibility with the pathogenic fibril architecture described here. The importance of Gly7 is further corroborated by its conservation in the human pathogenic proteins hSAA1.1 and hSAA1.3 (Fig. 4A) but not in mSAA2.1, mSAA3 and mSAA4, which carry a bulky, positively charged residue at this position and are unable to form amyloid *in vivo* (*9*).

**Fig. 4.**
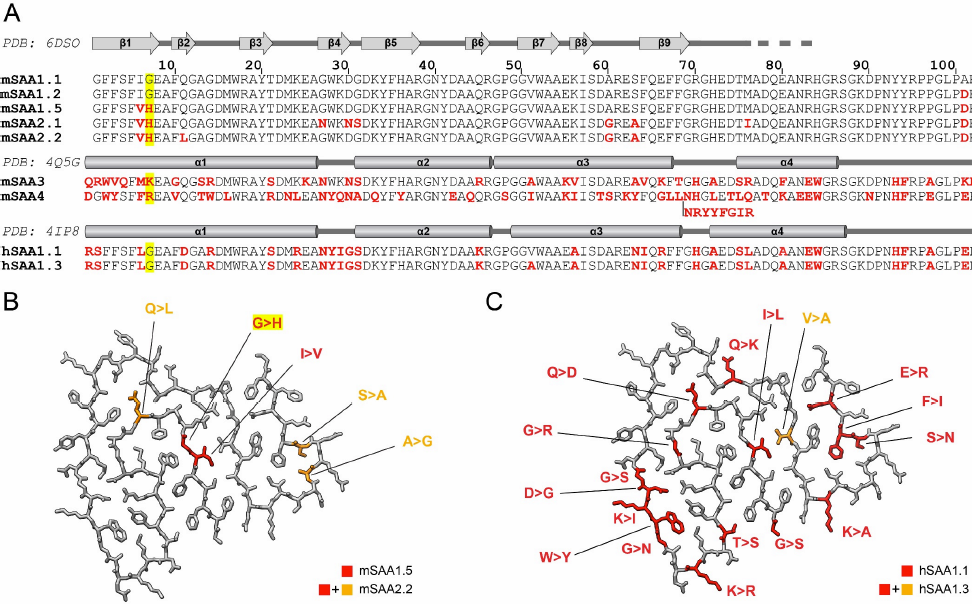
Comparison with naturally occurring SAA variants. (A) Sequence comparison. Red: amino acid substitutions compared to mSAA1.1. Secondary structural assignments according to the respective PDB entries are drawn above the sequence. Cylinders: α-helices; arrows: β-strands. (B) The substitutions in mSAA1.5 and mSAA2.2 are highlighted in the murine fibril structure. Both proteins additionally possess an Ala101Glu substitution, which does not confer amyloid resistance to mSAA1.2-expressing SJL/J mice (*30*). (C) The substitutions in hSAA1.1 and hSAA1.3 highlighted in the murine fibril structure.

Murine AA amyloidosis can be cross-seeded by amyloid fibrils purified from other species, including humans (*6, 26*). Although detailed AA amyloid fibril structures from other species are not known, the human fibril precursor proteins hSAA1.1 and hSAA1.3 lack substitutions within the structurally important hydrophobic core of the N-terminal β-arch (Fig. 4, A and C) and are thus compatible with this structural element. The β1-β7 zipper region shows only two conservative substitutions (Ile6Leu in both hSAA1.1 and hSAA1.3 and Val51Ala in hSAA1.3), while the fibril surface, C-terminal β-arch and the interface between N-terminal β-arch and intermediate region are affected by Gly14Arg, Asp30Gly, Glu66Arg and other non-conservative mutations, indicating conformational differences in these more C-terminal structural elements.

This first detailed structure of an amyloid fibril from systemic amyloidosis underscores the importance of working with *ex vivo* fibrils when drawing conclusions on the disease process. Our structure provides mechanistic clues about fibril formation and reveals a previously unknown relevance of electrostatic interactions for structuring AA amyloid fibrils. It explains the importance of the protein N-terminus for fibril formation and the resistance of two mouse variants to the development of amyloidosis. The observed protein fold and its compactness are remarkable given that mSAA1.1 was optimized by biological evolution to adopt a fundamentally different, but nevertheless compact conformation. The disease-associated fibril differs in morphology from all amyloid-like fibrils examined in this study and it is also more proteinase resistant (fig. S1). As proteinase resistance is a common feature of many *ex vivo* amyloid fibrils and prions (*14, 19, 27–29*), the ability to form a compact and/or protease resistant cross-β structure may explain the accumulation of only certain forms of aggregates inside the body. While clarifying these issues will require detailed structural information on other amyloid and amyloid-like fibrils, our current study demonstrates the general feasibility to obtain such information in systemic amyloidosis.

## Acknowledgments

The authors thank Paul Walther and Cornelia Loos (Ulm University) for technical support. This work was funded by the Deutsche Forschungsgemeinschaft (grant numbers FA 456/15-1 to M.F. and SCHM 3276/1 to M.S.). All cryo-EM data were collected at the European Molecular Biology Laboratory, Heidelberg (Germany), funded by iNEXT (Horizon 2020, European Union). F.L., S.L., M.R. A.S. and M.S. performed research. F.L., S.L., M.R. A.S., N.G., M.F. and M.S. analyzed data. P.W. and G.T.W. contributed tools and reagents. M.F. and M.S. designed research. F.L., S.L., N.G., M.F. and M.S. wrote the paper. None of the authors declares competing financial interest. The reconstructed cryo-EM map of the AA fibril has been deposited in the Electron Microscopy Data Bank with the accession code EMD-8910. The coordinates of the fitted atomic model have been deposited in the Protein Data Bank under the accession code 6DSO.

## Supplementary Materials for Cryo-EM structure of an amyloid fibril from systemic amyloidosis

**Authors:** Falk Liberta1#, Sarah Loerch^2#^, Matthies Rennegarbe^1^, Angelika Schierhorn^3^, Per Westermark^4^, Gunilla T. Westermark^5^, Nikolaus Grigorieff^2^, Marcus Fändrich^1^*, Matthias Schmidt^1^*.

^#^Both authors contributed equally.

^*^Correspondence to.

## Materials and Methods

### Recombinant mSAA proteins

Recombinant mSAA1.1, mSAA1.1(1-76) and mSAA1.1(1-82) were expressed and purified as described previously (*23*).

### Animal experiments and extraction of AA amyloid fibrils from murine tissue

Animal experiments were performed with female 6- to 8-week-old NMRI mice (Charles River Laboratories). Mice were divided into 4 experimental groups with two animals per group. On day 0, each mouse received a single 0.1 mL injection of a 0.1 mg/mL protein solution containing AA amyloid fibrils not treated with proteinase K (group A), AA amyloid fibrils treated with proteinase K (group B), non-fibrillar recombinant mSAA1.1(1-76) protein (group C) or a 1:1 mixture of recombinant, non-fibrillar mSAA1.1(1-76) and mSAA1.1(1-82) proteins (group D) into the lateral tail vein. Immediately afterwards each animal received a subcutaneous injection of 0.2 mL freshly prepared 1 % solution of AgNO3 in distilled water. The AgNO3 injection was repeated using 0.1 mL after 7 and 14 days. All animals were euthanized with CO2 on day 16. Spleens were removed, fixed in 4 % Roti‐Histofix formalin solution (Carl Roth) and embedded in paraffin for CR staining. AA fibrils from amyloid-laden mouse spleen were extracted as described previously (*13*). All experiments were approved by the Regierungspräsidium Tübingen.

### Formation of amyloid-like fibrils in vitro

Recombinant mSAA1.1 or mSAA1.1(1-76) were incubated at 1 mg/mL concentration in 50 mM sodium phosphate pH 3 or 10 mM tris(hydroxymethyl)aminomethane (Tris) pH 8 for 6 days at 37 °C and 300 rpm using a MTS2/4 digital microtiter shaker (IKA) placed in an BD53 incubator (Binder).

### Proteinase K treatment

A 180 μL aliquot from a stock solution of AA amyloid fibrils or amyloid-like fibrils at 0.3 mg/mL concentration was mixed with 20 μL 200 mM Tris buffer, pH 8.0, containing 1.4 M NaCl, 20 mM CaCl2. 0.4 μL proteinase K (20 mg/mL, Fermentas) were added to the mixture and incubated for 5 or 30 min at 37 °C. The proteinase K activity was stopped by adding 5 μL of 200 mM phenylmethane sulfonyl fluoride (Roth) prepared in methanol (Roth) to the reaction mixture and incubation for 10 min at room temperature.

### Amyloid grading of spleen section with Congo red

8-µm-thick spleen sections were cut with a HM 355S microtome (Thermo Fisher Scientific), deparaffinized with xylene for 20 min and rehydrated through a descending series of alcohol to distilled water (100 %, 90 %, 80 %, 70 %, 50 % and 0 % ethanol, 4 min each). Slides were stained with Hemalum solution acid according to Mayer (Carl Roth) for 1 min and rinsed with tap water for 15 min. Afterwards slides were stained with Congo red as described previously (*31*). Cover slips were mounted on microscopic slides using Roti‐Histokitt (Carl Roth) and samples were examined with an Eclipse 80i polarizing microscope (Nikon). Amyloid deposits were identified by characteristic green birefringence and graded as follows: 0 = no amyloid; 1 = isolated spots of amyloid within the tissue; 2 = perifollicular amyloid but rings not closed; 3 = closed rings of perifollicular amyloid; 4 = perifollicular amyloid rings and amyloid tissue infiltration.

### Denaturing gel electrophoresis

A solution of fibrils was mixed at 3:1 ratio with 4X lithium dodecyl sulfate (LDS) sample buffer (Thermo Fisher Scientific) and heated at 95 °C for 10 min. Proteins were separated on a NuPAGE 4–12 % Bis-Tris gradient gel (Thermo Fisher Scientific) using NuPAGE MES LDS running buffer (Thermo Fisher Scientific). The gel was stained for 1 h with a solution of 2.5 g/L Coomassie brilliant blue R250 in 20 % ethanol and 10 % acetic acid. The gel was destained in 30 % ethanol and 10 % acetic acid.

### Mass spectrometry

A sample of AA amyloid fibrils was dried by using a Vacuum Concentrator 5301 (Eppendorf) and resuspended in an equivalent volume of 6 M guanidine hydrochloride in 10 mM Tris buffer pH 8. The sample was desalted using a ZipTip (Merck Millipore). Matrix-assisted laser desorption/ionization MS spectra were recorded as described previously (*23*). Based on our set up a maximum error of 2 Da was assumed.

### Negative stain TEM

Negative-stain TEM specimens were prepared by loading 5 µL of the sample (0.2 mg/mL) onto a formvar and carbon coated 200 mesh copper grid (Plano). After incubation of the sample for 1 min at room temperature, the excess solvent was removed with filter paper. The grid was washed three times with water and stained three times with 2 % uranyl acetate solution. Grids were examined in a JEM-1400 transmission electron microscope (JEOL) that was operated at 120 kV.

### Platinum shadowing

Formvar and carbon coated 200 mesh copper grids (Plano) were glow-discharged using a PELCO easiGlow glow discharge cleaning system (TED PELLA). A 5 µl droplet of the AA amyloid fibril sample (0.2 mg/mL) were placed onto a grid and incubated for 30 s at room temperature. Excessive solution was removed with filter paper (Whatman). Grids were washed three times with water and dried at room temperature for 30 min. Platinum was evaporated at an angle of 30° using a Balzers TKR 010 to form a 1 nm thick layer on the sample. Grids were examined in a JEM-1400 transmission electron microscope (JEOL), operated at 120 kV.

### Cryo-EM

A 4 µL aliquot of a sample of AA amyloid fibrils (0.2 mg/mL) was applied to glow-discharged holey carbon coated grids (C-flat 2/1, 200 mesh), blotted with filter paper and plunge-frozen in liquid ethane using a Vitrobot Mark 3 (Thermo Fisher Scientific) Images were acquired using a K2-Summit detector (Gatan) in super-resolution counting mode on a Titan Krios transmission electron microscope (Thermo Fisher Scientific) at 300 kV. 1063 images that were composed of 40 frames each, were recorded in two sessions with exposure times of 0.6 s (session 1) and 0.5 s (session 2), exposure rates of 1.97 e^−^/Å^2^/s (session 1) and 2.52 e^−^/Å^2^/s (session 2) and a pixel size of 1.35 Å. Defocus values ranged from −1.3 to −5.5 µm.

### Helical reconstruction

Super-resolution movie frames were corrected for gain reference using IMOD (*32*). Motion correction, dose-weighting and binning by a factor of 2 was done using MOTIONCOR2 (*33*). To obtain a total electron dose of 20 e^−^/Å^2^ per aligned image, frames 3 – 18 (session 1) or frames 3 −8 (session 2) of each recorded movie were used. The contrast transfer function was estimated from the motion-corrected images using Gctf (*34*). Helical reconstruction was performed using RELION 2.1 (*35*). Fibrils were selected manually from the aligned micrographs. Segments were extracted using a box size of 210 pixel and an inter-box distance of 21 pixel (10 % of box length). Reference-free 2D classification with a regularization value of *T* = 2 was used to select class averages showing the helical repeat along the fibril axis. An initial 3D model was generated *de novo* from a small subset (200 particles per class) of the selected class averages using the Stochastic Gradient Descent algorithm implementation in RELION. The initial model was low-pass filtered to 20 Å and used for 3D auto-refinement to create a primary fibril model with an initial twist of 1.15° and 4.8 Å helical rise, as evident from the cross-over distance and the layer line profiles of the 2D classes. The resulting reconstruction showed clearly separated β-sheets (x-y plane) and partially resolved β-strands along the fibril axis. The generated primary model indicated the presence of two identical PFs, which are related by a pseudo-21 screw symmetry (4.8/2 Å rise and (360°-1.15°)/2 twist). Imposing this symmetry during reconstruction yielded a clear β-strand separation and side-chain densities. 3D classification (*T* = 4) with local optimization of helical twist and rise was used to further select particles for a final high-resolution auto-refinement. The best 3D classes were selected manually and reconstructed with local optimization of helical parameters using auto-refinement. All 3D classification and auto-refine processes were carried out using a central part of 10 % or 30 % of the intermediate asymmetrical reconstruction (*35*). The final reconstruction was post-processed with a soft-edge mask and an estimated map sharpening *B*-factor of −48 Å^2^. The resolution was estimated from the Fourier shell correlation (FSC) at 0.143 between the two independently refined half-maps.

### Model building, refinement and evaluation

An initial poly-alanine model was built *de novo* by using Coot (*36, 37*). Known geometries of -arches and arcades were considered for model building (*38, 39*). Once the backbone geometries were refined, side-chains were added. The clear densities for side-chains allowed us to unambiguously trace the orientation and register of the polypeptide chain. A PF fragment consisting of six subunits was assembled and refined with PHENIX (*40*) using phenix.real_space_refine (*41*) (phenix-1.13-2998). Non-crystallographic symmetry (NCS) restraints and constraints were imposed on all chains using a high-resolution cutoff of 3.2 Å. Initially, manually defined tight cross -sheet restraints were imposed and were relaxed at the later stages of refinement. Steric clashes, Ramachandran and rotamer outliers were detected during refinement using Molprobity (*42*) and iteratively corrected manually in Coot and refined in PHENIX. The PF fragment was then fitted into the density of the opposing PF. The final dodecamer was first refined using rigid body refinement where each protofilament was defined as one rigid body and finally using global minimization and atomic displacement parameter (ADP) refinement with secondary structure and NCS restraints until convergence. B-factor (ADP) refinement yielded a final B-factor of 79.7 Å (*37*). Detailed refinement statistics are shown in table S3.

The final model was evaluated using Molprobity. No Ramachandran outliers were detected and 97.01 % of the residues were in the favoured region of the Ramachandran plot. These were residues Ala44 and Gly49, which are found in kink regions of the fibril protein. An EMRinger score (*43*) of 6.10, calculated for the dodecamer, highlights excellent model-to-map fit at high resolution and high accuracy of backbone conformation and rotamers. -strands in the final model were analysed using DSSP (*44*) and STRIDE (*45*) and defined manually.

### Image representation

Image representations of reconstructed density (contoured at 1.5 σ) and refined model were created with UCSF Chimera (*46*) and PyMol. The following structures from the PDB were reproduced in the figures: mSAA3 (PDB 4Q5G) (*47*), hSAA1.1 (PDB 4IP8) (*48*), human phosphoglycerate kinase (PDB 3C39) (*49*). Final images were assembled with Adobe Illustrator CS6 (Adobe Systems).

**Fig. S1.**
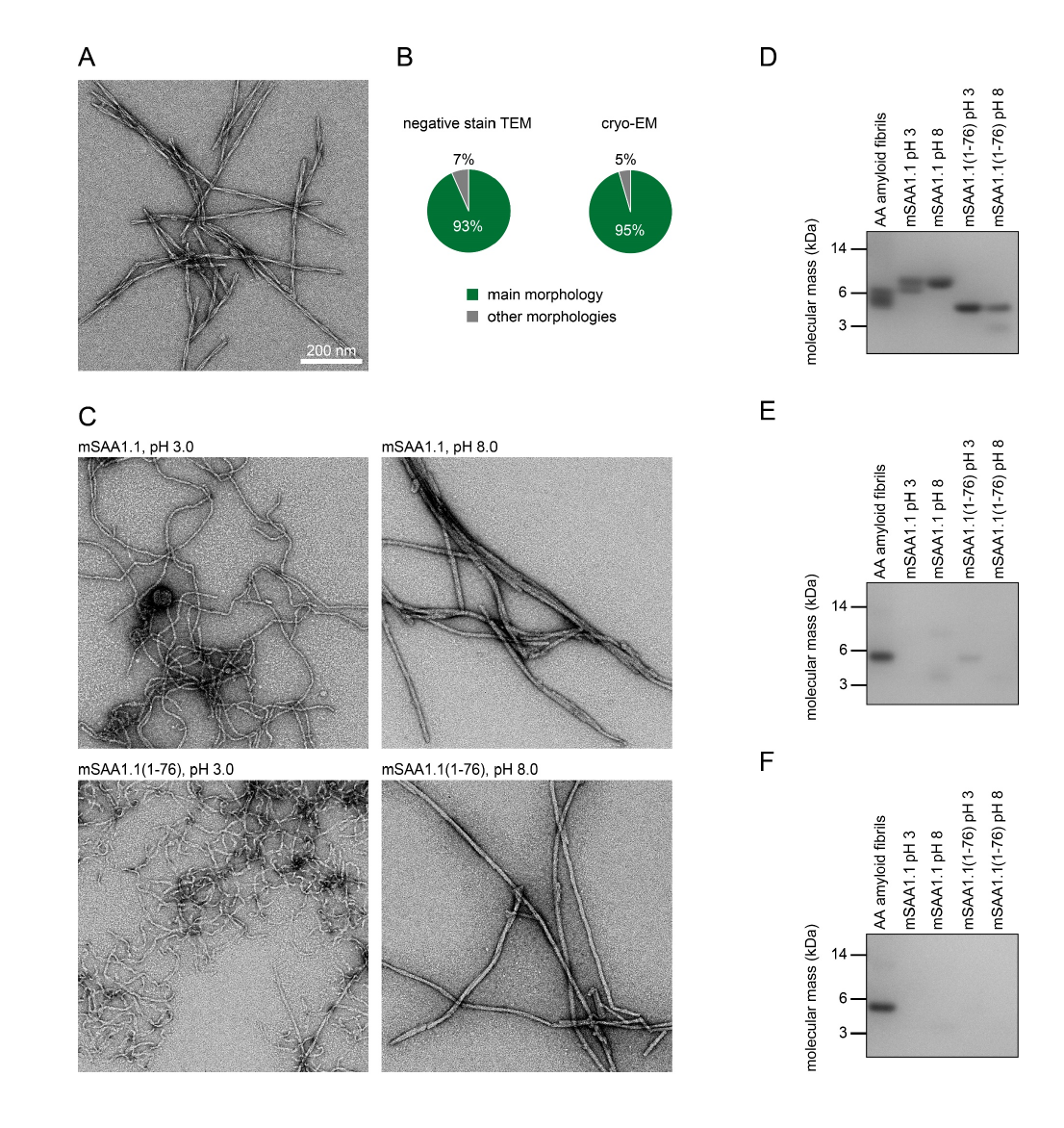
Structure of the extracted AA amyloid fibrils and of mSAA1.1-derived amyloid-like fibrils. (A) Negative stain TEM image of the extracted AA amyloid fibrils. (B) Quantification of the relative abundance of the main morphology as compared to other fibril morphologies based on negative stain TEM (n = 106 fibrils) and cryo-EM images (n = 220 fibrils). (C) Negative stain TEM image of the amyloid-like fibrils prepared *in vitro* from recombinant full-length mSAA1.1 protein and recombinant mSAA1.1(1-76). Two previously described conditions of *in vitro* fibril formation (*50*) were used (pH 3.0 and pH 8.0). The amyloid-like fibrils differ morphologically from the AA amyloid fibrils. (D-F) Denaturing gel electrophoresis of fibrils shown in (A) and (C). (D) Samples prior to proteinase K treatment. (E) Fibrils exposed to proteinase K for 5 min. (F) Fibrils exposed to proteinase K for 30 min.

**Fig. S2.**
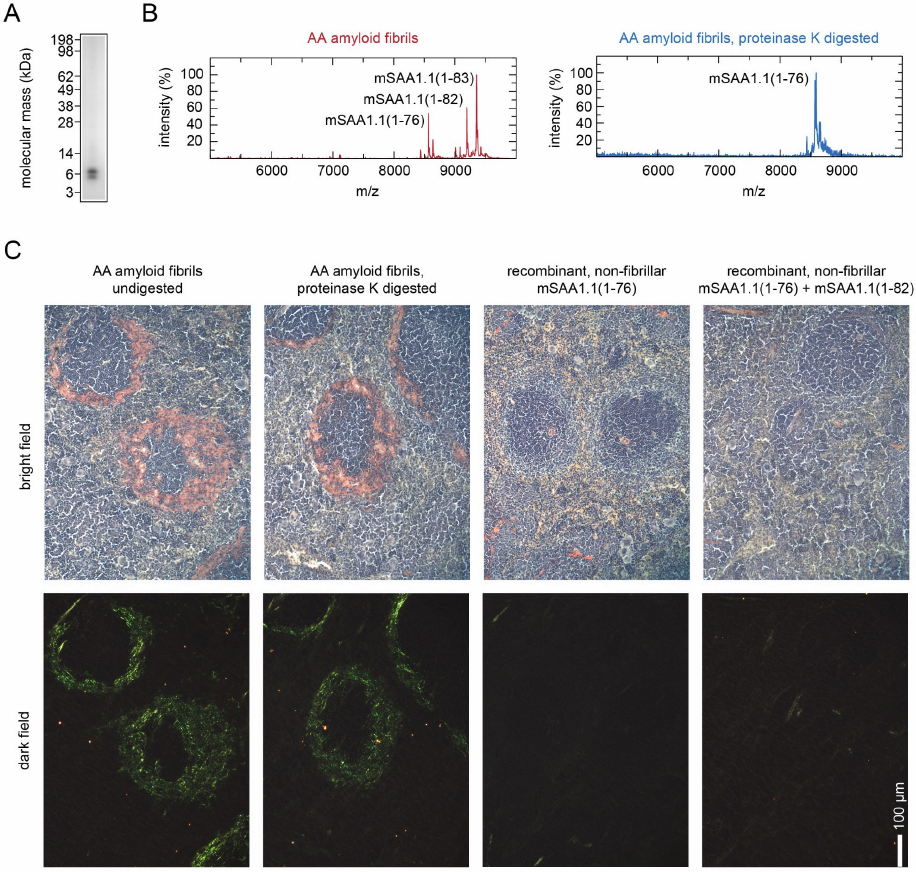
Molecular composition and prion-like activity of AA amyloid fibrils. (A) Coomassie-stained denaturing protein gel prepared with AA amyloid fibrils. (B) Mass spectra obtained with AA amyloid fibrils before (left) and after proteinase K treatment (right). Observation of the mSAA1.1(1-76) fragment after proteinase K treatment is consistent with the known cleavage specificity of proteinase K (*51*); that is, the C-terminus of Met76 represents the first proteinase K cleavage target beyond residue 69. (C) Representative bright field and dark field polarizing microscopy images of Congo red stained section of murine spleen. Mice were injected with AA amyloid fibrils without and after proteinase K treatment. The amyloid grade is +3 in both cases. Mice receiving injections of non-fibrillar samples consisting of freshly dissolved, recombinant mSAA1.1(1-76) or of a 1:1 mixture of dissolved, recombinant mSAA1.1(1-76) and mSAA1.1(1-82) did not develop amyloid (amyloid grade 0).

**Fig. S3.**
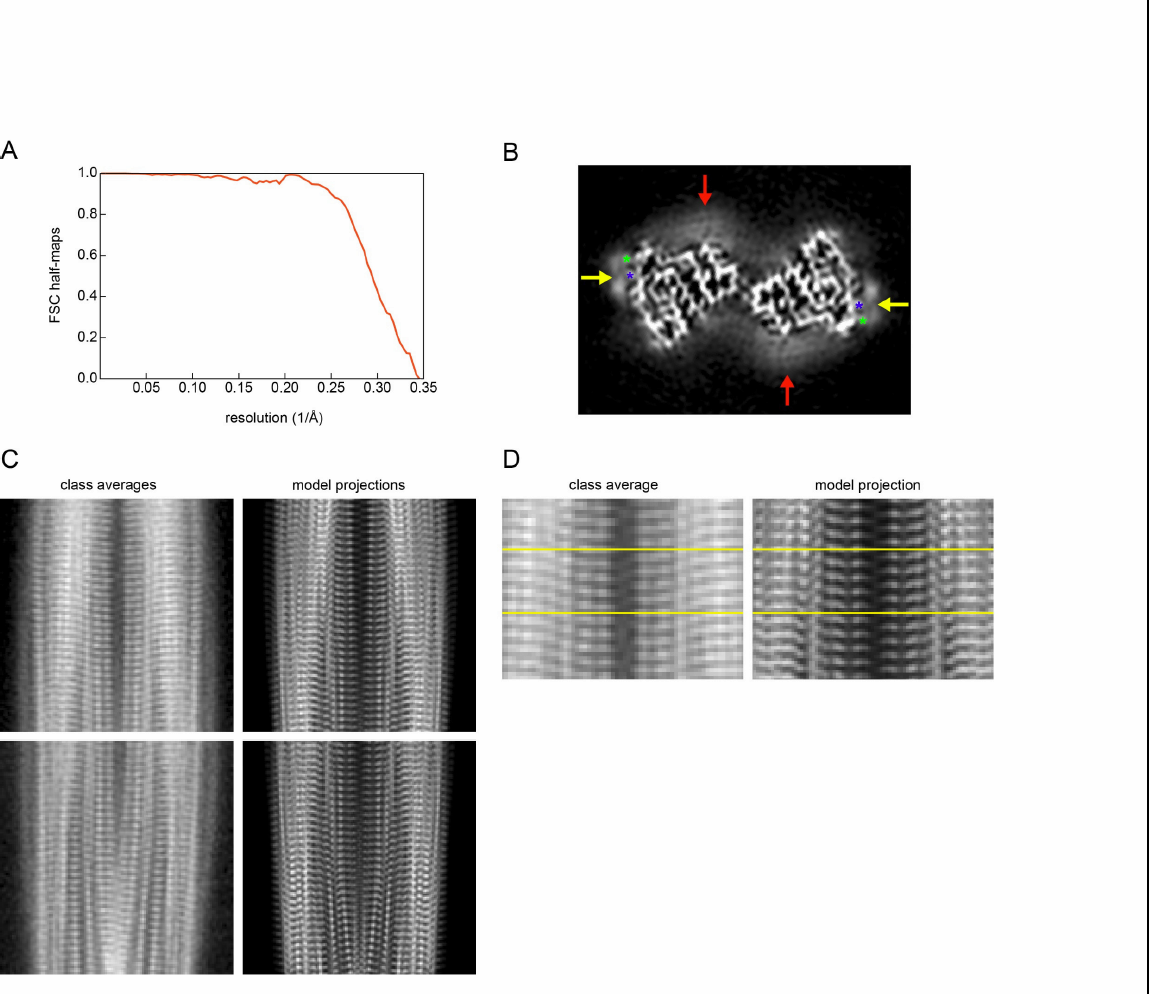
Details of the reconstructed 3D map. (A) FSC between the two half maps. (B) A 6.75 Å thick cross-sectional slice of the unmasked 3D map. Red arrow: weak density adjacent to the C-terminus, indicating structural disorder at residues 70-83. Yellow arrow: weak density adjacent to a small basic surface patch formed by His36 (blue star) and Arg38 (green star) that were previously implicated in glycosaminoglycan catalyzed fibril formation (*52*). (C) Comparison of representative 2D class averages and corresponding 2D projections obtained with the model. (D) Close-up of a 2D class average and corresponding 2D projection to highlight the stagger of the two PFs. Yellow lines are drawn to guide the eye.

**Fig. S4.**
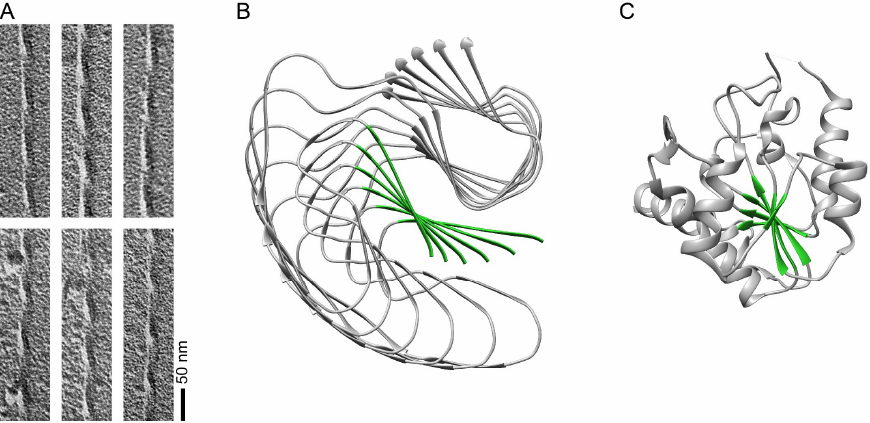
Twisted structure of AA amyloid fibrils. (A) Transmission TEM images of AA amyloid fibrils after platinum side shadowing. (B) Left-hand twist illustrated by the β1 sheet (green) of the fibril model. Only every tenth protein chain along the fibril axis is displayed. (C) Left-hand β-sheet twist in residues 1-202 of human phosphoglycerate kinase (PDB 3C39). The left-hand twist is defined by the counter-clockwise rotation (front to back) when viewed along the backbone hydrogen bonds or in the direction of the fibril axis.

**Fig. S5.**
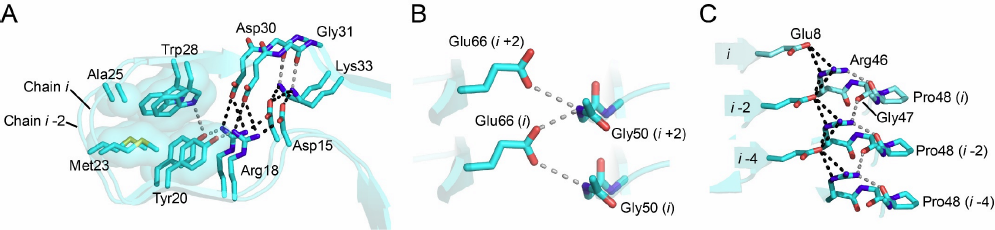
Buried ionic and polar interactions. (A) Intramolecular salt-bridges are formed between Asp15, Arg18 (both β3), Asp30 (β4) and Lys33 (β5). Lys33 additionally forms an intermolecular salt-bridge with Asp15 of chains *i* and *i*-2. The boundary with the adjacent hydrophobic patch is formed by Tyr20 (β3) and Trp28 (β4), which form hydrogen bonds with Arg18 and with each other on one side, while packing with their respective hydrophobic surfaces against Met23 (β3-β4 arc) on the other side. (B) Glu66 (β9) forms bidentate electrostatic interactions with the backbone nitrogens of Gly50 of chains *i* and *i*+2. (D) Arg46 (β6) forms electrostatic interactions with the carbonyl oxygen of Pro48 (β6-β7 arc), and intra- and intermolecular salt bridges with Glu8 (β1) in chains *i* and *i*-2. Salt bridges: back; hydrogen bonds: grey.

**Fig. S6.**
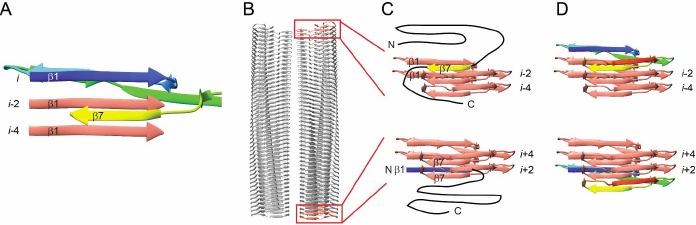
Possible structure-based mechanism for the fibril replication. (A) Packing of the intermolecular zipper between strand β7 (molecule *i*) and strands β1 (molecules *i*-2 and *i*-4). (B) Side view of a fibril section with the top and bottom two molecules of the right PF highlighted. (C, D) Scheme for the extension of the amyloid fibril. (C) Addition of the strand β7 (top) or β1 (bottom) from a new molecule helps to orient the incoming polypeptide chain at the fibril tip. (D) Once assembled, the added polypeptide chain exposes a β1 (top) or β7 (bottom) half zipper itself. The exposure of different half zippers at either end of the polar fibril implies different mechanisms and/or outgrowth kinetics to occur at the two fibril ends, which was observed with other fibril systems (*15, 53*).

**Table S1.** **Molecular species detected by MS before and after proteinase K treatment.** Left column: peak maximum position detected by MS; central column: assigned protein species; right column: calculated protein mass.

| Peak position (m/z) | Assigned protein fragment | Theoretical mass (Da) |
| --- | --- | --- |
| ***Before proteolysis*** |  |  |
| 8564.2 | mSAA1.1(1-76) | 8565.2 |
| 9194.4 | mSAA1.1(1-82) | 9193.8 |
| 9349.5 | mSAA1.1(1-83) | 9350.0 |
| ***After proteolysis*** |  |  |
| 8565.1 | mSAA1.1(1-76) | 8565.2 |

**Table S2.** Structural statistics of cryo-EM data collection and image processing.

| ***Data Collection*** |  |
| --- | --- |
| Microscope | Titan Krios (Thermo Fisher Scientific) |
| Camera | K2 Summit (Gatan) |
| Acceleration voltage (kV) | 300 |
| Magnification | x 105,000 |
| Defocus range ($\mu\text{m}$) | -1.3 to -5.5 |
| Dose rate ($\text{e}^-/\text{\AA}^2/\text{s}$) | 1.97 and 2.52 |
| Number of movie frames | 17 and 16 |
| Exposure time (s) | 10.2 and 8 |
| Total electron dose ($\text{e}^-/\text{\AA}^2$) | 20 |
| Pixel size ($\text{\AA}$) | 1.35 |
| ***Reconstruction*** |  |
| Box size (pixel) | 210 |
| Inter box distance ($\text{\AA}$) | 28 |
| Number of extracted segments | 137,956 |
| Number of segments after 2D classification | 93,045 |
| Number of segments after 3D classification | 21,024 |
| Resolution based on the 0.143 FSC criterion ($\text{\AA}$) | 3.0 |
| Map sharpening B-Factor ($\text{\AA}^2$) | -48 |
| Helical rise ($\text{\AA}$) | 2.41 |
| Helical twist ($^\circ$) | 179.44 |

**Table S3.** Structural statistics of model building and refinement.

|  |  |
| --- | --- |
| ***Model composition*** |  |
| Non-hydrogen atoms | 6588 |
| Number of chains | 12 |
| ***Model Refinement*** |  |
| Resolution | 3.3 |
| Map CC (around atoms) | 0.7836 |
| RMSD bonds (Å) | 0.009 |
| RMSD angles (°) | 0.928 |
| All-atom clash score | 1.53 |
| Ramachandran outliers/favoured (%) | 0/97.01 |
| Rotamer outliers | 0.00 |
| C-beta deviations | 0 |
| EMRinger score | 6.10 |
| EMRinger Zscore | 13.86 |
| Molprobity score | 1.07 |

